# *ARID1A* Mutations Protect Follicular Lymphoma from FAS-dependent Immune Surveillance by Reducing RUNX3/ETS1-Driven FAS-Expression

**DOI:** 10.1101/2023.12.12.571212

**Authors:** Martina Antoniolli, Maria Solovey, Carolin Dorothea Strobl, Deepak Bararia, William David Keay, Johannes Adrian Hildebrand, Louisa Adolph, Michael Heide, Verena Passerini, Tabea Freyholdt, Lucas Wange, Wolfgang Enard, Susanne Thieme, Helmut Blum, Martina Rudelius, Julia Mergner, Christina Ludwig, Sebastian Bultmann, Marc Schmidt-Supprian, Heinrich Leonhardt, Marion Subklewe, Michael von Bergwelt-Baildon, Maria Colomé-Tatché, Oliver Weigert

## Abstract

The cell death receptor FAS and its ligand (FASLG) play crucial roles in the selection of B cells during the germinal center (GC) reaction. Failure to eliminate potentially harmful B cells via FAS can lead to lymphoproliferation and the development B cell malignancies. The classic form of follicular lymphoma (FL) is a prototypic GC-derived B cell malignancy, characterized by the t(14;18) (q32;q21)IGH::*BCL2* translocation and overexpression of antiapoptotic BCL2. Additional alterations were shown to be clinically relevant, including mutations in *ARID1A*. ARID1A is part of the SWI/SNF nucleosome remodeling complex that regulates DNA accessibility (“openness”). However, the mechanism how *ARID1A* mutations contribute to FL pathogenesis remains unclear.

We analyzed 151 FL biopsies of patients with advanced stage disease at initial diagnosis and found that *ARID1A* mutations were recurrent and mainly disruptive, with an overall frequency of 18%. Additionally, we observed that *ARID1A* mutant FL showed significantly lower FAS protein expression in the FL tumor cell population. Functional experiments in BCL2-translocated lymphoma cells demonstrated that ARID1A is directly involved in the regulation of FAS, and ARID1A loss leads to decreased FAS protein and gene expression. However, ARID1A loss did not affect *FAS* promotor openness. Instead, we identified and experimentally validated a previously unknown co-transcriptional complex consisting of RUNX3 and ETS1 that regulates *FAS* expression, and ARID1A loss leads to reduced *RUNX3* promotor openness and gene expression. The reduced FAS levels induced by ARID1A loss rendered lymphoma cells resistant to both soluble and T cell membrane-anchored FASLG-induced apoptosis.

In summary, we have identified a functionally and clinically relevant mechanism how FL cells can escape FAS-dependent immune surveillance, which may also impact the efficacy of T cell-based therapies, including bispecific antibodies and CAR T cells.

## Introduction

Avoiding immune destruction is a hallmark of cancer (1). This is particularly prominent in malignant lymphomas, the most common type of blood cancer (2). Most B cell Non-Hodgkin lymphomas (B-NHL) originate from germinal center (GC) B cells. Germinal centers (GC) are highly specialized microstructures within lymphoid tissues where GC reactions occur, involving a complex interplay among B cells, T cells, and antigen-presenting cells. During the GC reaction, B cells undergo iterative rounds of genetic mutations of their immunoglobulin genes followed by selection to ultimately produce higher affinity immunoglobulins (3). The cell death receptor FAS and its ligand (FASLG) play crucial roles in the germinal center reactions. Normal GC B cells express high levels of FAS and low levels of antiapoptotic BCL2 (4), rendering them prone to apoptosis induction. Failure to eliminate potentially harmful B cells can lead to accumulation of abnormal B cells and-eventually-the development of B-NHLs (and/or autoimmunity).

Follicular lymphoma (FL) is a prototypic GC-derived B-NHL and a clinically and molecularly highly heterogeneous disease. The molecular hallmark of classic FL is the translocation t(14;18)(q32;q21) IGH::*BCL2* (5, 6), which is acquired in early B cells (7) and leads to aberrant overexpression of BCL2. Yet, the *BCL2* translocation alone is insufficient for lymphomagenesis. Additional genetic and epigenetic alterations contribute to the development of FL and regulate critical interactions with the tumor microenvironment (TME) (8).

Many groups including us are increasingly untangling the multifaceted genetic landscape of FL (9, 10). We have previously shown that distinct gene mutations are linked with the clinical course and treatment outcome in patients with advanced stage FL receiving standard immunochemotherapies (11, 12), including mutations in *ARID1A*. ARID1A is part of a multimeric SWItch/Sucrose Non-Fermentable (SWI/SNF) nucleosome remodeling complex which plays a pivotal role in regulating chromatin structure (13). By altering DNA accessibility (“openness”) it is involved in the regulation of gene expression (14–16). *ARID1A* is recurrently mutated in FL at initial diagnosis (10, 11) with an increase from <10% in biopsies from limited stage disease to >20% in advanced stage FL (17). These mutations are mostly heterocygous and disruptive, leading to protein haplodeficiency (18). However, the contribution of ARID1A loss to FL development and progression and to the biology of the disease remains unclear.

Interestingly, a previous functional genome-wide shRNA screen had shown that knock-down of *ARID1A* rescued a variety of cancer cell lines from FASLG-induced apoptosis (19). Therefore, and because of the described interrelationships, we decided to study the link between ARID1A loss and FAS/FASLG-induced apoptosis in FL. Here, we show that *ARID1A* mutations disrupt a previously unknown regulatory network controlling FAS expression that involves RUNX3 and ETS1 and promotes a functionally and clinically relevant immune evasive phenotype in FL.

## Results

### *ARID1A* mutations are associated with reduced FAS levels in human FL biopsies

First, we wanted to test the hypothesis that *ARID1A* mutations are associated with lower FAS levels in human FL. For this, we re-analyzed our previously reported cohort of diagnostic biopsies from patients with advanced stage FL at initial diagnosis (11), consisting of 151 evaluable cases with available targeted DNA sequencing data that included *ARID1A* gene mutation status. Thereof, 51 cases had also been analyzed by digital multiplex gene expression profiling (DMGEP) that included *FAS* expression levels (20), and 43 cases were available for quantitative multispectral imaging (QMI) that included staining for FAS (**Fig. 1A**).

**Figure 1:**
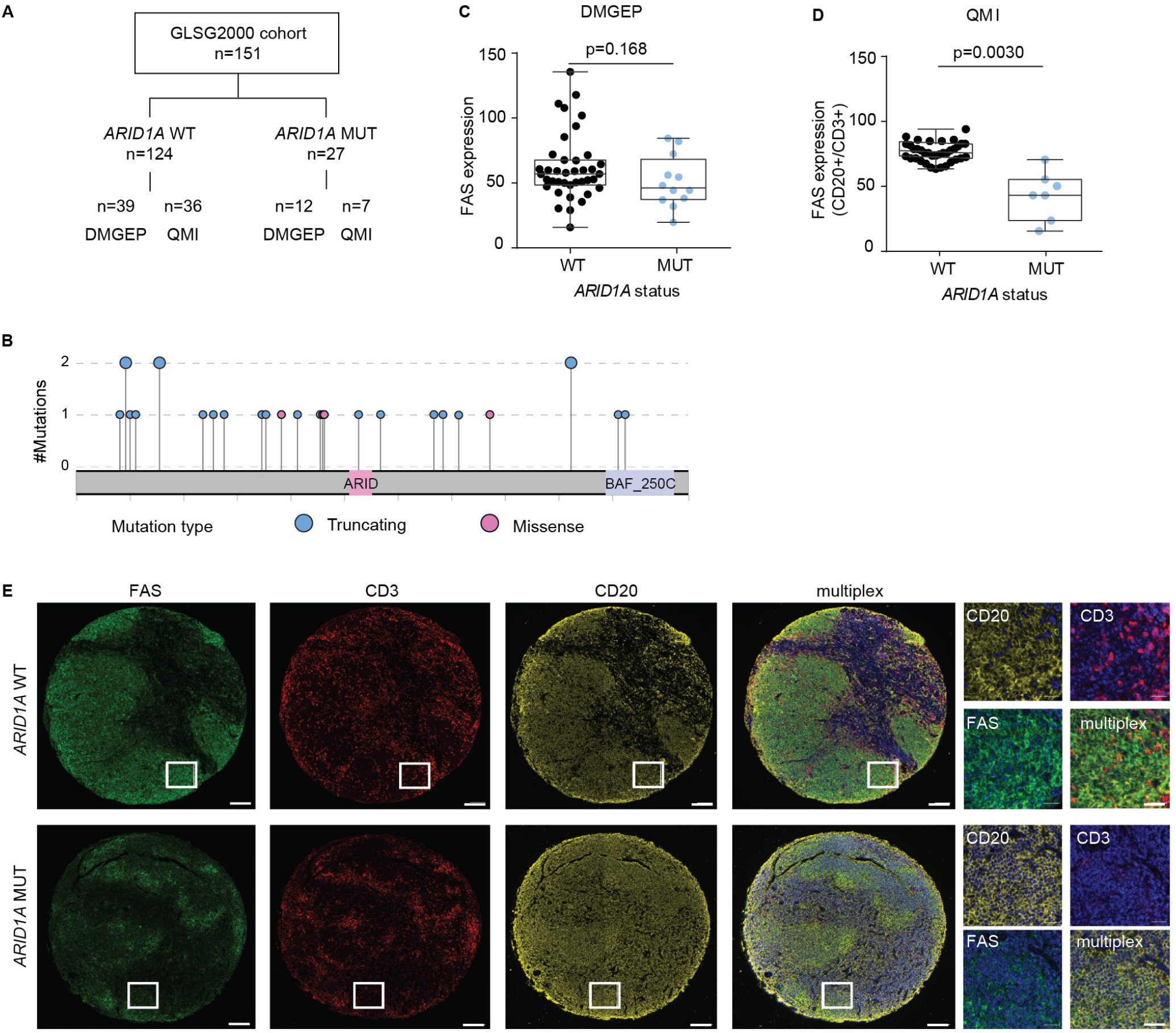
*ARID1A* mutations are associated with low FAS levels in primary human FL biopsies. **A** Schematic overview of the GLSG2000 FL cohort and available data. **B** Lollipop plot of *ARID1A* mutations in the evaluable GLSG2000 FL cohort. **C** *FAS* RNA expression in primary FL biopsies (*ARID1A*^WT^ (N = 39) *vs ARID1A*^MUT^ (N = 12)) by digital multiplex gene expression profiling (DMGEP). *P-values from Mann-Whitney U-test.* **D** FAS protein abundance in the CD20^+^ cells normalized to CD3^+^ cells in primary FL biopsies (*ARID1A*^WT^ (N = 36) *vs ARID1A*^MUT^ (N = 7)) by quantitative multispectral imaging (QMI). *P-values from Welch test.* **E** Representative multispectral images. Scale bar is 20 µm (low magnification) or 400 µm (high magnification).

The overall frequency of non-synonymous *ARID1A* mutations (at allelic fractions 5%) was 18% (27/151), the majority categorized as disruptive and distributed throughout the coding region of the gene (23/27, 88%) (**Fig 1B**). *ARID1A* mutant (*ARID1A*^MUT^) FL (evaluable N = 12) showed a trend towards lower overall *FAS* gene expression levels compared to *ARID1A* wild type (*ARID1A*^WT^) cases (N = 39) by DMGEP (**Fig 1C**). As DMGEP represents bulk gene expression data and *FAS* is known to be highly expressed by non-FL cells such as T cells and macrophages of the TME we performed QMI, which allows single cell resolution. Quantification of FAS protein expression in the FL tumor cell population revealed lower expression in *ARID1A*^MUT^ (N = 7) *vs ARID1A*^WT^ (N = 36) FL (**Fig 1D-E**). Overall, this data shows that *ARID1A* mutations are highly recurrent, predominantly disruptive and associated with lower FAS expression in advanced stage primary human FL.

### *ARID1A* disruption results in decreased FAS protein expression

For mechanistic and functional studies, we studied human B cell lymphoma cell lines that harbor the FL-hallmark t(14;18)(q32:q21)IGH::*BCL2* translocation, including two *ARID1A*^WT^ cell lines (OCI-Ly1 and OCI-Ly8) and two *ARID1A*^MUT^ cell lines (Karpas422 and WSU-FSCCL). We confirmed lower ARID1A expression in *ARID1A*^MUT^ cells by Western blot (**Suppl Fig 1A**) and flow cytometry showed corresponding lower FAS expression on *ARID1A*^MUT^ cell lines compared to *ARID1A*^WT^ cell lines (**Suppl Fig 1B**). To demonstrate that ARID1A is directly involved in the regulation of FAS expression, we introduced heterocygous (het) or homocygous (hom) *ARID1A* deletions into the *ARID1A*^WT^ cell lines OCI-Ly1 and OCI-Ly8 by CRISPR/Cas9 and generated single-cell derived clones. Immunoblotting confirmed lower ARID1A expression (*i.e.*, haplodeficiency) in *ARID1A*^het^ cells, and complete knock-out (KO) in *ARID1A*^hom^ cells (**Fig 2A**). Next, we evaluated FAS cell surface expression on these cells by flow cytometry. Again, we observed reduced FAS levels on *ARID1A*^het^ and KO cells (**Fig 2B**). Of note, ectopic re-expression of ARID1A in *ARID1A*^het^ cells (OCI-Ly8) restored FAS levels (**Fig 2B**), indicating that ARID1A is directly involved in the regulation of FAS expression. We validated lower FAS protein expression using targeted proteomics: FAS protein levels were lower in *ARID1A*^het^ and KO cells, both in cell surface proteins as well as in the total proteome fraction, and could be rescued by re-expression of ARID1A, respectively **Fig 2C**).

**Figure 2:**
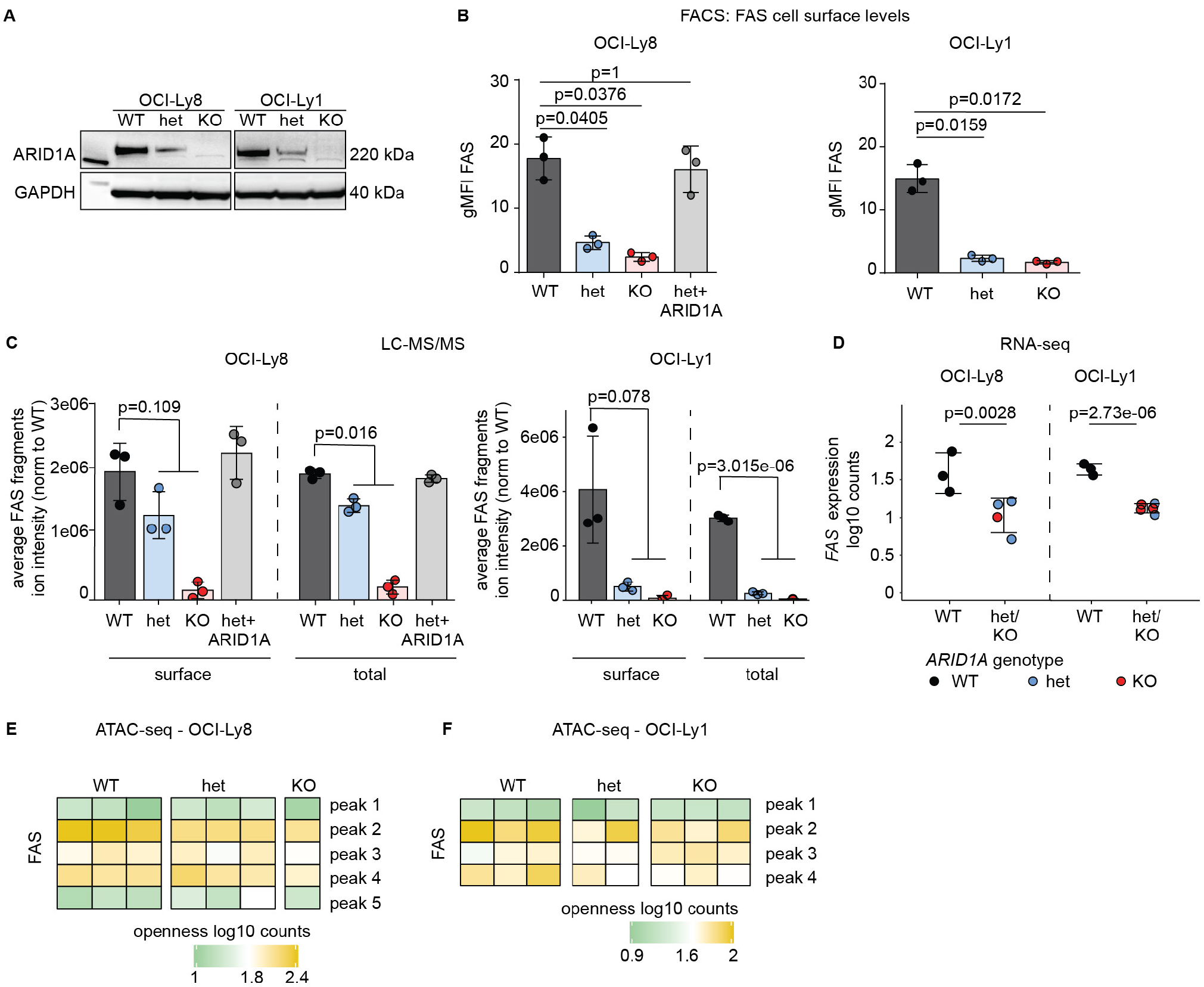
ARID1A loss results in decreased *FAS* gene expression but does not affect FAS promotor openness. **A** Western blot of single-cell derived clones of OCI-Ly8 and OCI-Ly1 cells with *ARID1A*^WT^ (WT) or CRISPR-Cas9-introduced heterocygous *ARID1A* mutation (het) or *ARID1A* knockout (KO). **B** FAS cell surface expression on OCI-Ly8 (left panel) and OCI-Ly1 (right panel) clones by FACS. Bar diagram for the geometric means of independent biological replicates (N = 3). *P-values for OCI-Ly8 are from two-sided t-test, OCI-Ly1 from Welch test, Bonferroni adjusted. Each group was tested against WT.* **C** FAS peptide abundance by targeted proteomics in the surface proteome and total proteome lysates of OCI-Ly8 (left panel) and OCI-Ly1 (right panel) clones (N = 3). *P-values from two-sided t-test were used. Het and KO values were tested together against WT.* **D** *FAS* RNA expression by RNA-Seq in OCI-Ly8 and OCI-Ly1 single-cell-derived clones. *P-values from the DESeq2 R package for differential expression analysis with the default Benjamini-Hochberg correction were used. Het and KO values were tested together against WT.* **E** *FAS* promotor accessibility (five detected peaks) measured by ATAC-Seq in OCI-Ly8 clones. **F** *FAS* promotor accessibility (four detected peaks) measured by ATAC-Seq in OCI-Ly1 clones. *Pooled data from biological replicates (N) are represented as mean± SD*.

### ARID1A loss results in decreased *FAS* gene expression but does not affect *FAS* promotor openness

Next, we tested whether *FAS* gene expression levels were affected by ARID1A loss. Indeed, both RNA sequencing and quantitative real-time PCR (qRT-PCR) showed lower *FAS* gene expression levels in *ARID1A*^het^ and KO cells (**Fig 2D** and **Suppl Fig 1C**).

We hypothesized that ARID1A regulates FAS protein levels by directly affecting *FAS* promotor chromatin accessibility and performed Assay for Transposase-Accessible Chromatin using Sequencing (ATAC-Seq). Differential promotor openness analysis revealed 241 and 206 differentially open (DO) peaks for OCI-Ly1 and OCI-Ly8, respectively (**Table S1-2**). However, we did not detect differences in chromatin accessibility at the *FAS* promotor upon ARID1A loss (**Fig 2E** and **2F**).

### Identification of the *FAS*-regulating RUNX3/ETS1 co-transcriptional complex

Next, we hypothesized that the lower FAS levels upon ARID1A loss may be explained by altered expression of FAS-regulating transcription factors (TFs) or co-transcription factors (co-TFs) (**Fig 3A**). For this, we first used the DoRothEA database (21–23) to identify all TFs which are directly involved in the regulation of *FAS* (**Table S3**). We tested the differential expression of the TFs directly involved in *FAS* regulation in *ARID1A*^MUT^ *vs ARID1A*^WT^ cells in both OCI-Ly1 and OCI-Ly8 (**Fig. 3A**, “Hypothesis I”). However, we could not identify mutual differentially expressed *FAS*-regulating TFs that correlated with *ARID1A* mutation status/loss (**Fig 3A**). Thus, the lower FAS levels upon ARID1A loss cannot be explained by differential expression of a direct *FAS*-regulating TF. We then turned into testing the differential expression of their co-TFs (**Fig. 3A**, “Hypothesis II”). These co-TFs were identified as having been shown or predicted to physically interact with these *FAS*-regulating TFs, utilizing the STRING database (24) of protein-protein interactions (**Fig 3A, Table S4**). In both cell lines, we identified *RUNX3* to be differentially expressed upon ARID1A loss and predicted to be an interaction partner of ETS1, which has previously been found to bind to the *FAS* promotor (25, 26). Accordingly, the promotor openness and RNA expression of *ETS1* were unchanged (**Fig 3B, C,** and **Suppl Fig 2 A, C**), whereas the promotor of *RUNX3* was partially closed and its RNA expression was reduced upon ARID1A loss (**Fig 3B, D**, **Table S2**, and **Suppl Fig 2B, D, E**). This suggested that ARID1A loss leads to reduced *FAS* expression through reduced *RUNX3* promotor openness and reduced expression of *RUNX3*, which interacts with ETS1, a putative *FAS*-regulating TF.

**Figure 3:**
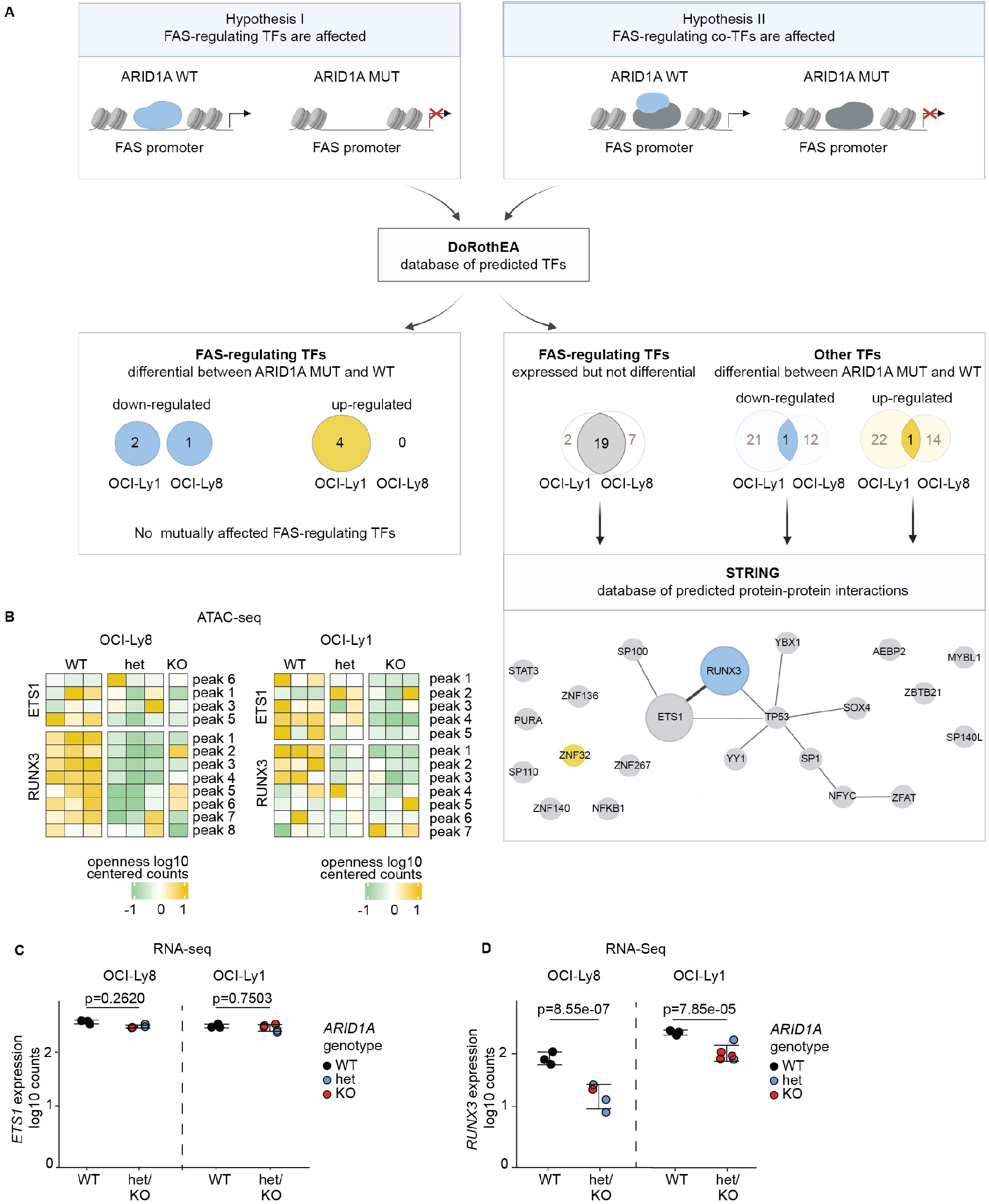
Identification of the *FAS*-regulating RUNX3/ETS1 co-transcriptional complex. **A** Analysis workflow and results of FAS-regulating transcription factors (TFs) and co-transcription factors (co-TFs). **B** ATAC-Seq differential openness analysis of *ARID1A*^MUT^ single-cell-derived clones *vs ARID1A*^WT^ control clones, in OCI-Ly8 (left panel) and OCI-Ly1 (right panel). Heatmap of centered log10 counts of all detected (open) peaks in the promotor regions of *ETS1* and *RUNX3*. **C** *ETS1* gene expression analysis by RNA-Seq in *ARID1A*^MUT^ clones (blue for het, red for KO) *vs ARID1A*^WT^ clones (black). **D** *RUNX3* gene expression analysis by RNA-Seq in *ARID1A*^MUT^ clones (blue for het, red for KO) *vs ARID1A*^WT^ clones (black). *P-values for panels C and D were calculated using the DESeq2 package for differential expression analysis with the default Benjamini-Hochberg correction. Het and KO values were tested together against WT. Pooled data from biological replicates (N) are* _29_ *represented as mean*± *SD*.

### Experimental validation of the *FAS*-regulating RUNX3/ETS1 co-transcriptional complex

To functionally validate this novel FAS-regulatory network, we first analyzed the genomic context around the *FAS* promotor (**Fig 4A**) and searched for ETS1 binding motifs (**Fig 4B**). We identified putative ETS1 binding sites that mapped to the *FAS* promotor region and overlapped with open peaks in our ATAC sequencing data as well as with previously reported ETS1 binding sites (25). Then, we cloned two different sized fragments (537 bp and 332 bp) from that region into a luciferase reporter construct. The reporter constructs were expressed in HEK 293T cells, along with increasing doses of ETS1 or RUNX3 (**Suppl Fig 3A**). We could show dose-dependent transactivation activity for ETS1 (**Fig 4C**) confirming that ETS1 itself is a direct *FAS*-transactivating TF. In contrast, RUNX3 expression alone did not show transactivation activity (**Suppl Fig 3B**).

**Figure 4:**
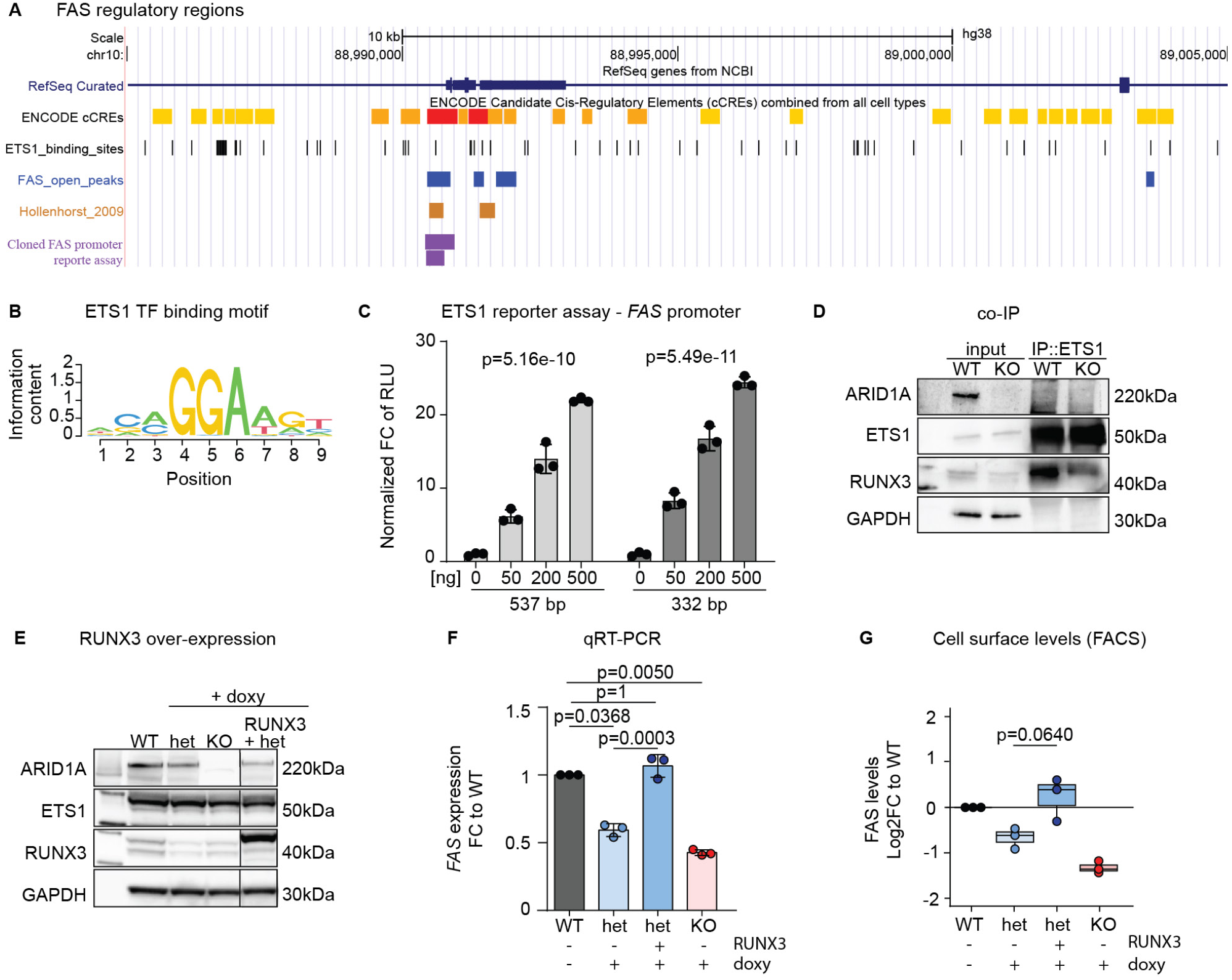
Experimental validation of the *FAS*-regulating RUNX3/ETS1 co-transcriptional complex. **A** Schematic of the *FAS* gene with annotated enhancer regions (yellow) and promotor (red), ETS1 binding sites (black), accessible chromatin regions from our ATAC-Seq data (blue), ETS1 binding sites from published ChIP-Seq data (brown) (24, 25), and the *FAS* promotor regions cloned for the reporter assay (purple). **B** ETS1 TF binding motif in *FAS* accessible promotor regions. **C** Luciferase reporter assay with co-transfection of the ETS1 expression vector and pGL3-FAS constructs in 293T cells (N = 3). *P-values are from linear regression model on square root transformed dose and non-transformed response values.* **D** Western blot of inputs and ETS1-immunoprecipitated OCI-Ly1 (WT and KO). **E** Western blot of OCI-Ly8 clones (*ARID1A* WT, het, and KO) with or without stable doxycycline (dox)-induced overexpression of RUNX3. **F** Rescue of *FAS* RNA levels upon RUNX3 overexpression in OCI-Ly8 measured by quantitative real-time PCR (TaqMan assay) (N = 3). *P-values are from two-sided t-test, Bonferroni adjusted. All groups were tested against WT, and het+RUNX3 was tested against het.* **G** Rescue of FAS cell-surface protein levels upon RUNX3 over expression in OCI-Ly8 measured by FACS (N = 3). *P-value is from two-sided t-test. Het+RUNX3 was tested against het. Pooled data from biological replicates (N) are represented as mean ±SD*.

Next, we wanted to test whether ETS1 and RUNX3 are direct interaction partners. For this, we performed co-immunoprecipitation in OCI-Ly1 cells, i.e. pull-down of ETS1 and immunoblotting for RUNX3. As shown in **Fig 4D**, we could detect direct interaction of ETS1 and RUNX3. Of note, RUNX3 abundance was lower in cells with ARID1A loss (KO), both in the input and the pull-down sample (**Fig. 4D**). Furthermore, we confirmed that ARID1A loss (het and KO) had no impact on ETS1 protein levels in these cells, but RUNX3 levels were reduced (**Fig 4E**).

Finally, we over-expressed RUNX3 in cells with ARID1A loss (het) (**Fig 4E, Suppl Fig 3C**). Immunoblotting confirmed high expression of RUNX3 while ETS1 levels were not affected (**Fig 4E, Suppl Fig 3C**). RUNX3 overexpression indeed resulted in increased *FAS* expression upon RUNX3 overexpression, both on the transcriptional level as shown by qRT-PCR (**Fig 4F, Suppl Fig 3D**) as well as on the protein level as shown by flow cytometry (**Fig 4G, Suppl Fig 3E**). In summary, these experiments validate our model of an ARID1A-dependent RUNX3/ETS1-mediated network regulating *FAS* expression.

### ARID1A loss leads to functionally relevant reduction of FAS/FASLG-induced apoptosis

Finally, we wanted to test whether reduced FAS expression upon ARID1A loss is functionally relevant. Upon binding of FAS ligand (FASLG), FAS oligomerizes and forms the death-inducing signaling complex (DISC), activating the extrinsic apoptotic pathway (27). Thus, we treated cells with or without ARID1A disruption with increasing doses of purified soluble human recombinant FAS ligand (FASLG) and assessed apoptosis by flow cytometry (**Fig 5A**). In both lines, clones with ARID1A loss were less sensitive to FASLG treatment (**Fig 5B**). Of note, RUNX3 overexpression restored sensitivity towards FASLG treatment (**Fig 5B**).

**Figure 5:**
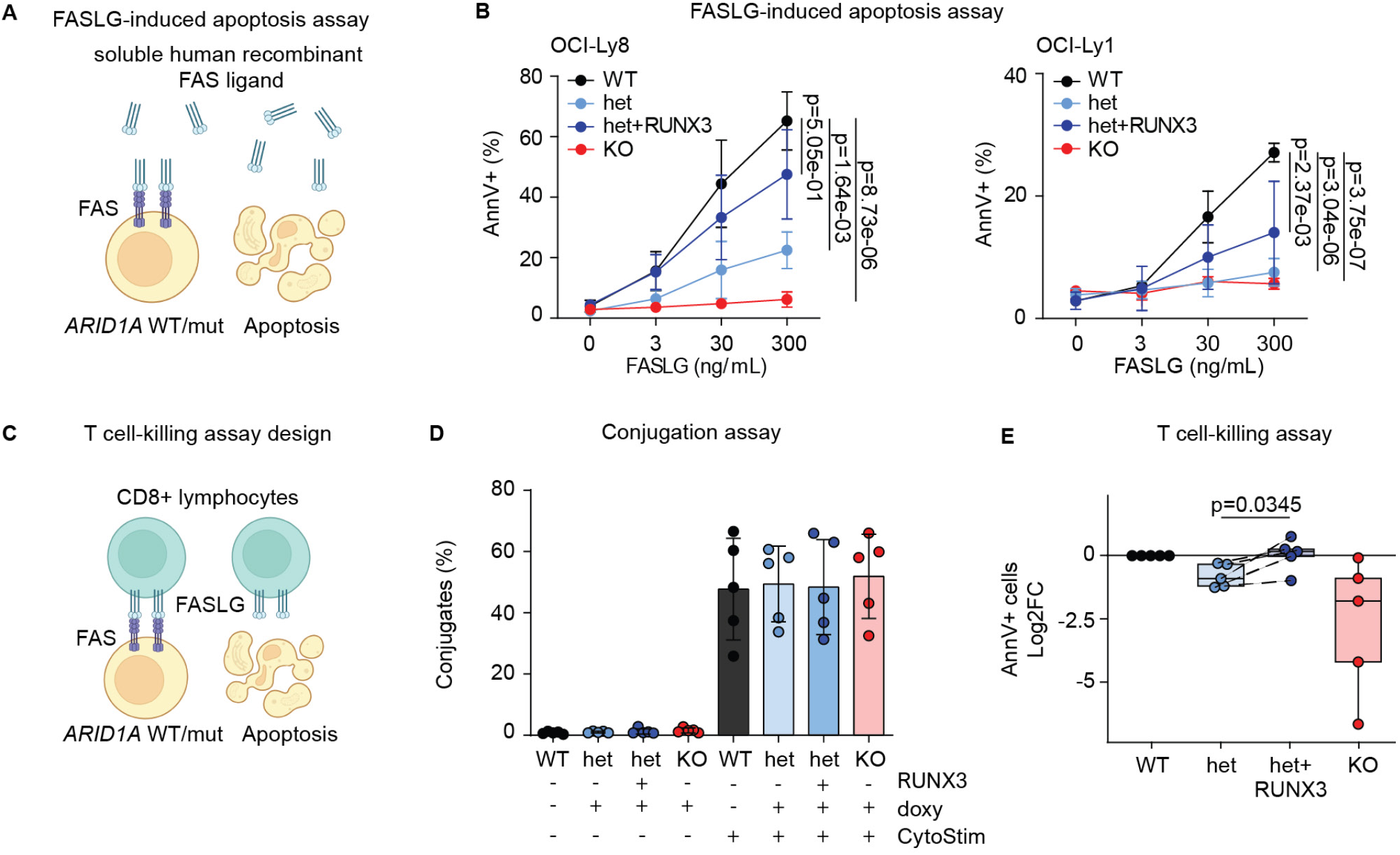
ARID1A loss leads to functionally relevant reduction of FAS/FASLG-induced apoptosis. **A** Schematic of the FASLG-induced apoptosis assay (soluble human recombinant FAS ligand treatment). **B** Percent AnnexinV-positive OCI-Ly8 (left panel) and OCI-Ly1 (right panel) clones with *ARID1A* WT, het, KO and overexpression of RUNX3 on het (het + RUNX3) after 24 h treatment with increasing dose of purified soluble human recombinant FAS ligand (N = 3). *P-values are from a linear regression on square root transformed values tested against WT and Bonferroni-adjusted*. **C** Schematic of the T-cell mediated killing assay. **D** Conjugate formation of OCI-Ly8 cells (CFSE^+^) and CD8^+^ T-cells (VPD^+^); double-positive conjugates (CSFE^+^/VPD^+^) quantified as percentage of total CD8^+^ T-cells (VPD^+^) (N = 5) at indicated conditions. **E** T cell mediated cytotoxicity; percentage of OCI-Ly8 cells (CFSE^+^) undergoing apoptosis upon co-culture with CD8^+^ T cells quantified as percentage of all measured cells at indicated conditions (N = 5, biological replicates performed with different healthy T cell donors). *P-value is from the paired Welch test. Het+RUNX3 was tested against het. Pooled data from biological replicates (N) are represented as mean ±SD*.

Normally, FAS is mostly engaged by membrane-anchored FASLG on the surface of cytotoxic cells, particularly activated T cells and NK cells (28). Therefore, we co-cultured CFSE-labelled lymphoma cells (OCI-Ly8) with or without ARID1A loss (*ARID1A*^het^ or ARID1A^KO^ *vs ARID1A*^WT^ clones) with VDP-labelled T cells (CD8^+^) from five different healthy donors (**Fig 5C**). While conjugate formation of lymphoma cells and T cells were not affected by *ARID1A* genotype (**Fig 5D**), cells with ARID1A loss were less sensitive to T cell-mediated apoptosis, and overexpression of RUNX3 restored sensitivity towards T cell-mediated killing (**Fig 5E**).

Overall, these experiments demonstrate that ARID1A loss leads to functionally relevant reduced FAS levels in lymphoma cells, rendering cells resistant to both soluble and membrane-bound FASLG-induced apoptosis.

## Discussion

*ARID1A* is among the ten most commonly mutated genes in cancer (29, 30). Its pleiotropic effects in different settings, its mutation pattern and distribution across a wide variety of different tumor entities as well as apparently conflicting functional and clinical data support the concept that ARID1A functions as a context-dependent tumor suppressor (31). This highlights the importance of investigating its function in specific tumor types and settings.

Here, we show that ARID1A loss results in a biologically and clinically relevant immune evasive phenotype in FL by rendering tumor cells resistant to FASLG-induced apoptosis. Specifically, we identify and functionally characterize a novel *FAS*-regulating network, which involves reduced RUNX3 and ETS1-driven *FAS* expression upon ARID1A loss (**Fig 6**), which may have therapeutical implications, as discussed below.

**Figure 6:**
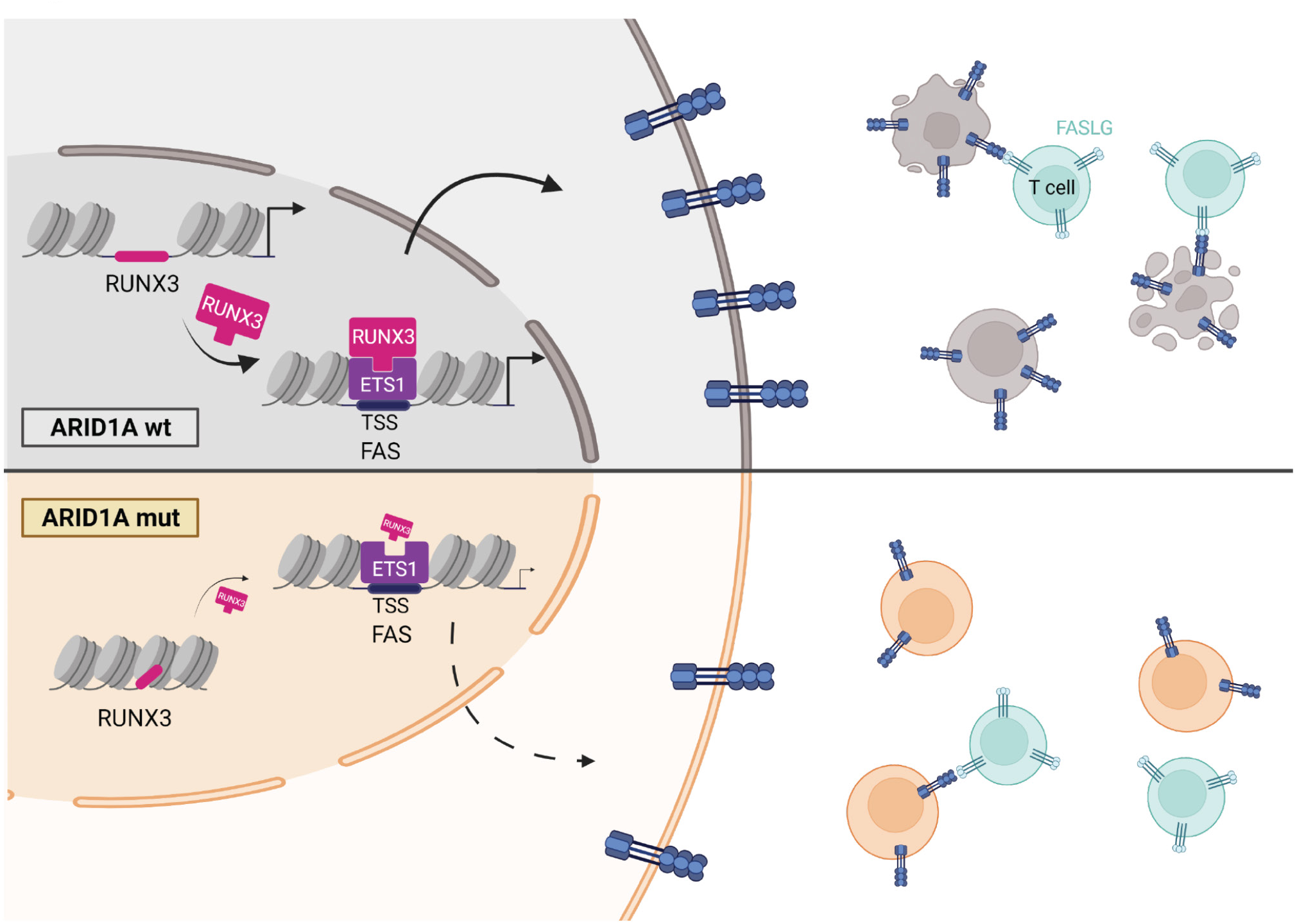
Graphical abstract. Low FAS expression in *ARID1A* mutant lymphoma cells is mediated by reducing RUNX3/ETS1-driven FAS transcription and promotes a functionally and potentially clinically relevant escape from T cell mediated killing.

We initially identified *ARID1A* mutations in FL more than 10 years ago, when we reported a rare case of donor-derived FLs. Following a bone marrow transplantation, both the donor and the recipient developed FLs that originated from a common precursor clone (18). Interestingly, both FLs harbored distinct *ARID1A* mutations that each resulted in protein haplodeficiency, *i.e.* these mutations were acquired individually (18). This convergent evolution already suggested that ARID1A loss provides a selective advantage during the development and progression of FL. Now, our functional data sheds light on the underlying biology. GC T cells express both FASLG and CD40L and the decision to kill or help largely depends on the expression of their receptors on B cells (32). *ARID1A* mutant (pre-) malignant B cells with low FAS are more likely to evade FASLG-mediated negative selection during the GC passage, which explains the high mutation frequency in FL at initial diagnosis (10, 11). Similarly, *ARID1A* mutant (sub-) clones may evade T cell immunosurveillance during disease progression or relapse, which would explain the additional accumulation of *ARID1A* mutations in advanced stage and relapsed/refractory (r/r) FL (17)).

In fact, accumulation of *ARID1A* mutations in r/r FL may profoundly impact the efficacy of subsequent therapies, maybe most notably of chimeric antigen receptor (CAR) T cell therapies. CAR T cells targeting CD19 have demonstrated high response rates patients with r/r FL (33, 34). Yet, the majority of patients will ultimately relapse and the mechanisms of treatment failure are an area of active research. There is evolving preclinical and clinical evidence that FAS loss is associated with CAR T cell failure. *E.g.*, a genome-wide CRISPR/Cas9 screen identified the necessary role for FAS/FASLG in CD19-directed CAR T cell killing in an *in vivo* model of B cell lymphoma (35). Importantly, this study also showed that low tumoral FAS expression was predictive of poor outcome in patients with relapsed or refractory large B cell lymphoma (LBCL) in the pivotal ZUMA-1 trial of axicabtagene ciloleucel (axi-cel) (35). Moreover, focal deletion of 10q23.3 leading to FAS loss was found to be associated with shorter progression-free survival (PFS) and overall survival (OS) in 122 evaluable patients who received CD19-directed CAR T cells for r/r LBCL (36). Thus, we hypothesize that other alterations that lead to clinically and functionally relevant FAS loss, including mutations in *ARID1A* as described herein and in *FAS* itself, will be enriched in patients who fail CAR T cell therapies.

Vice versa, killing of tumor cells by T cells activated / recruited by bispecific monoclonal antibodies (BiAbs) has been shown to depend mostly on perforin and granzyme A/B activity rather than FAS/ FASLG (37, 38). This may explain the sustained and remarkable activity of BiAbs even in patients who failed CAR T cell therapies, as shown for example for mosunetuzumab, a CD20 × CD3 T cell-engaging BiAbs in patients with r/r FL (39). Ultimately, future studies will clarify whether *ARID1A* mutations and/or other alterations that result in FAS loss can serve as clinically useful biomarkers to guide optimal sequencing of therapies for patients with r/r FL.

Finally, our functional data demonstrates that the FAS phenotype in cells with ARID1A loss is mediated through reduced accessibility and transcription of the *FAS*-regulating co-TF *RUNX3*. Yet, RUNX3 itself may have tumor suppressive function. For example, 45%-60% of human gastric cancers do not express RUNX3 due to hemicygous deletion and hypermethylation of the *RUNX3* promotor, and *Runx3* knock-out mice develop hyperplasia of the gastric mucosa (40). Of note, *RUNX3* is located (in close proximity to *ARID1A*) on the distal portion of the short arm of human chromosome 1 (1p36), which is commonly deleted in FL (41, 42). Thus, RUNX3 may be a *bona fide* tumor suppressor in FL and be inactivated by several mechanisms, including 1p36 deletions and/or disruptive *ARID1A* mutations. Last but not least, *ARID1A* mutations as well as loss of RUNX3 may have other functional consequences in addition to reducing FAS levels in FL that could be of clinical relevance and this should be investigated further.

In summary, we have elucidated the molecular mechanism of reduced FAS expression in *ARID1A* mutant lymphoma cells and demonstrated that this promotes functionally and potentially clinically relevant escape from T cell mediated killing.

## Materials and methods

### Primary patient samples

Gene mutation data and gene expression data from diagnostic biopsies were derived from our previously reported cohorts of patients with previously untreated advanced stage FL (11, 20). All studies on human material were covered by IRB approvals (LMU #223-14, LMU #276-14, LMU #056/00, and LMU #539-15 fed).

### Multispectral imaging analysis (Vectra® Polaris System)

Multiplex immunohistochemistry was performed as described previously (20). We used the following antibodies: FAS-R (EP208; AC-0178RUO; Abcam, Cambridge, UK; 1:100), CD3 (SP7; RBK024-05; Zytomed System, Berlin, Germany; 1:150); CD20 (L26; 120M-85; Cell Marque, Rocklin, CA, USA; 1:200). Images were acquired using the quantitative slide scanner with the Vectra® Polaris 1.0. (PerkinElmer, Waltham, MA, USA) and analyzed using InForm 2.4.2 (PerkinElmer) and HALO® software (Indica Labs, Albuquerque, NM, USA). Antibody details are listed in **Table S6**.

### Cell lines

Karpas422 and WSU-FSCCL carry heterocygous *ARID1A* mutations (Q1959fs and copy number loss (43), respectively) and were cultured in RPMI 1640 (PAN^TM^ Biotech, Aidenbach, Germany). OCI-Ly1 and OCI-Ly8 are *ARID1A* wild type and were cultured in IMDM (PAN^TM^ Biotech), each supplemented with 10% heat-inactivated FBS (PAN^TM^ Biotech). All cell lines were authenticated by STR profiling (Eurofins, Val Fleuri, Luxembourg) and were tested negative for mycoplasma (MycoAlert PLUS mycoplasma detection kit, Lonza, Basel, Switzerland).

### CRISPR/Cas9-mediated knockdown of ARID1A

Single guide RNA (sgRNA) targeting *ARID1A* (**Table S5**) were cloned into the pSpCas9(BB)-2A-GFP backbone (PX458, Addgene plasmid #48138) as previously described (44). OCI-Ly8 or OCI-Ly1 cells were transduced with Nucleofector^TM^ Solution V (Lonza) and the Nucleofector^TM^2b (Lonza) and single-cell sorted for GFP after 48 hours.

### Construction of lentiviral vectors and cell lines

Overexpression of ARID1A was performed by lentiviral transduction (pHAGE-CMV-MCS-ARID1Awt-IHRES-ZsGREEN) as previously described (20). To express *RUNX3* we used the lentiviral construct pTet-O-RUNX3-T2A-PuroR (Addgene # 162349).

### Flow Cytometry

Cells washed with PBS (PAN) and stained with anti-CD95 (FAS) antibody for 45 min at 4°C (PE anti-human CD95 (FAS) mouse DX2 (# 305608) BioLegend (San Diego, CA, USA) (1:25)). Cells were washed and resuspended in 200 µL PBS for FACS analysis. At least 10,000 events were recorded. Antibody details are listed in **Table S6**.

### Quantitative real-time PCR analysis

Total RNA was isolated using Direct-zol^TM^ RNA Kits (Zymo Research, Irvine, CA, USA) and transcribed into cDNA using iScript cDNA Synthesis Kit (BioRad, Hercules, CA, USA). PCR reactions were performed using TaqMan^TM^ Fast Advanced Master Mix, FAS (Hs00236330_m1 FAM-MGB FAS), RUNX3 (Hs01091094_m1 FAM-MGB RUNX3), and TBP (Hs00427620_m1 VIC-MGB TBP) assays (Thermo Fisher Scientific). The statistical analysis was done on RT-qPCR cycle threshold (ct) values normalized to housekeeping gene.

### Western Blot

Cells were lysed with radio immunoprecipitation assay buffer (RIPA). Protein concentrations were quantitated with Pierce BCA assay (Thermo Fisher Scientific). Proteins were separated on 4-12% SDS-PAGE gels. Primary antibody incubation was performed overnight at 4°C, followed by a secondary horseradish peroxidase-conjugated antibody at room temperature for 1 h. The following antibodies were used: ARID1A, rabbit (#HPA005456), Sigma Aldrich (St. Louis, MO, USA) (1:2500); RUNX3, mouse, clone R3-5GA (#697901), BioLegend (1:2000); ETS1, rabbit, clone D8O8A (#14069S), Cell Signaling (1:2000); GAPDH, mouse, clone 6C5 (#MA5-15738-D680), Thermo Fisher Scientific (1:10000). Antibody details are listed in **Table S6**. Uncropped Western blots are provided in Supplementary Material.

### Co-immunoprecipitation (IP)

Cells were lysed using Pierce^TM^ IP Lysis Buffer (Thermo Fisher Scientific) supplemented with protease and phosphatase inhibitor cocktails (Roche, Basel, Switzerland). IP was performed with 3 mg of protein. SureBeads^TM^ Protein A Magnetic Beads (BioRad) were coupled with anti-ETS1 antibody (CS#14069, Cell Signaling, Danvers, MA, USA) for 3 h at 4°C with continuous rotation. Lysates were incubated with the bead-bound antibody overnight at 4°C with continuous rotation. Bead-bound immunoprecipitates were washed three times with IP buffer and eluted twice with a total volume of 2x Laemmli Buffer (BioRad). Input samples (30 µg) and co-IP samples were subjects to western blot analysis. Antibodies are listed in **Table S6**. Uncropped Western blots are provided in Supplementary Material.

### Mass Spectrometry

Cell surface proteins were isolated by using Pierce Cell Surface Protein Isolation Kit (Thermo Fisher Scientific, cat#89881), following the manufacturers’ instructions. The bound proteins were eluted with 350 µl Novex^TM^ Tris-Glycine Native Sample Buffer (2X) (Thermo Fisher Scientific), supplemented with DTT to a final concentration of 50 mM. In-gel trypsin digestion was performed according to standard procedures (45).

For total proteome analysis, cells were lysed in 2% SDS and 50 mM Tris-HCl pH 7.5 and heated to 95°C for 5 min. 1 μL 100 TFA was added to each sample to hydrolyze DNA and the pH subsequently adjusted to 8.5 with 3 M Tris solution. SP3 cleanup was performed according to the protocol by Hughes *et al.* (46) followed by tryptic digestion overnight and stage-tip desalting. Dry peptides were reconstituted in 2% (v/v) acetonitrile, 0.1% (v/v) formic acid in HPLC grade water and spiked with PROCAL retention time standard peptide mix (47).

Liquid chromatography-coupled mass spectrometry (LC-MS/MS) analysis was performed on a Q Exactive HF-X Orbitrap (Thermo Fisher Scientific) coupled on-line to a Dionex Ultimate 3000 RSLCnano system (Thermo Fisher Scientific). FAS protein abundance was monitored using a parallel reaction monitoring assay (PRM) with a 50 min linear gradient. The recorded RAW files were imported into Skyline (64-bit, v.20.2.0.343) for data filtering and analysis. For FAS protein detection the six most intense and unique FAS peptides were selected. A spectral library was constructed using the PROSIT prediction algorithm implemented in Skyline with standard settings (48, 49). Peaks were integrated using the automatic settings followed by manual curation of all peak boundaries. Peaks with a dotp product < 0.7 compared to the predicted peptide spectrum were excluded from analysis. For FAS protein quantification the area of all fragment ion traces over all peptides was summed.

### FAS ligand (FASLG)-induced apoptosis assay

*ARID1A* wild-type and mutant cell lines were treated with 0, 3, 30, or 300 ng/mL purified soluble human recombinant FAS ligand (SUPERFASLIGAND®, Enzo Life Sciences, Inc., Farmingdale, New York, USA) for 24 h. The cells were assayed by flow cytometry (BD FACSCanto™ II). We utilized the AnnexinV Apoptosis Detection Kit I (BD PharmingenTM, San Diego, CA, USA) and DAPI (BD PharmingenTM). The data was analyzed with the FlowJo v10 software (BD PharmingenTM).

### Co-culture assays

Lymphoma cells were stained with 1 μM CellTrace^TM^ CFSE Cell Proliferation Dye (Thermo Fisher Scientific, Waltham, Massachusetts, USA) for 4 min at room temperature (RT). CD8^+^ T cells were isolated from human peripheral blood using EasySep^TM^ Human CD8^+^ T Cell Isolation Kit (Stemcell Technologies, Vancouver, BC, Canada) and stained with 10 μM CellTrace^TM^ Violet Cell Proliferation Kit (VPD) (Thermo Fisher Scientific) for 10 min at 37°C. CytoStim^TM^ (Miltenyi Biotec, Cologne, Germany) was used according to the manufacturers’ recommendations to activate T cells by binding the T cell receptor (TCR) and crosslinking it to major histocompatibility complex (MHC) molecules of B cells. Cells were cultured in TexMACS media (Miltenyi Biotech) in a 1:1 ratio at 37°C up to 3 h, stained with AnnexinV and assayed by FACS as described above. Conjugate formation was calculated as [CFSE^+^/VPD^+^ cells VPD^+^ cells] 100%. The specific CD8^+^ T cell-mediated cytotoxity (T cell-killing assay) was calculated by subtracting the unspecific AnnexinV^+^ lymphoma cells (*i.e.*, cells cultured without T cells as a control) from AnnexinV^+^ CFSE-labelled lymphoma cells co-cultured with CD8^+^ T cells, and quantified as a percentage of all measured cells as previously described (50).

### Luciferase Reporter Assay

Two regions of *FAS* promoter (537bp and 332bp) were cloned into pGL3-basic vector following the manufacturer’s instruction (#E1751, Promega, Walldorf, Germany). The cloned regions (FASprom_P1 and FASprom_P2, **Table S5**) contained ETS1 TF binding motifs (25). The cells were analyzed by Dual-Glo® Luciferase Assay (Promega #E2920).

### RNA-Sequencing

RNA-sequencing was performed on 8 or 7 different clones for OCI-Ly1 and OCI-Ly8, respectively, each clone being sequenced in two technical replicates. RNA was isolated using Direct-zol^TM^ RNA MicroPrep kit (Zymo Research) and a total of 10 ng of RNA per sample were subjected to RNA sequencing using a version of the prime-seq protocol (51) Illumina paired end sequencing was performed on an HiSeq 1500 (Illumina, San Diego, CA, USA) instrument. The first 16 bp read was used for sample barcode and UMI, the second 50 bp read was used for the gene. Raw data was demultiplexed using deML (52) and processed using the zUMIs pipeline (53) with STAR (54). Reads were mapped to the human genome (hg38) with Ensemble gene annotations (GRCh38.84) and annotated using biomaRt (v2.44.4). Genes with the mean raw counts less or equal to 5 counts were filtered. Technical replicates were collapsed using the collapseReplicates() function of the DESeq2 package (v1.28.1). Counts were normalized using DESeq2 package (v1.28.1). Differentially expressed genes were calculated with DESeq2 package (v1.28.1) between the controls and mutants (het + KO). Only genes that had log2 fold change higher than 1 or lower than −1, as well as the adjusted p-value lower than 0.1 were considered as significantly differentially expressed (**Table S7**).

A total of fifteen single-cell-derived clones were assayed in OCI-Ly1 and OCI-Ly8. Control clones (3 per cell line) were transduced with the empty pSpCas9(BB)-2A-GFP backbone. In OCI-Ly1, two heterocygous and three ARID1A knock-out clones were sequenced. In OCI-Ly8, three heterocygous and one knock-out clone were sequenced.

### ATAC-Sequencing

Transposased fragments were prepared and pre-amplified as described in the optimized Omni-ATAC protocol (55). Transposased DNA was amplified using Nextera XT Index pair (i5 Index Name, Illumina), specific for every sample (list in **Table S8**), and the KAPA HiFi PCR Kit (Roche Diagnostics), as recommended by the manufacturer. Amplified fragments were purified and eluted. Illumina paired end sequencing was performed on an HiSeq 1500 instrument, where the first 16 bp read covered the sample barcode and UMI, and the second 50 bp read was used to identify the gene. Reads were trimmed using TrimGalore (v0.6.5; default parameters), aligned to the reference genome (hg38) with Bowtie2 (v2.3.5; --very-sensitive -X 2000) and sorted by position and indexed using samtools (v1.2). Those mapping to mitochondria, mapping to less than 6 bases or with a MAPQ below 10 were removed. Alignments of fragments longer than 150bp were also filtered using Deeptools-alignmentSieve (v3.3.1). Peak calling was done using MACS2-callpeak (v2.2.6; --nomodel -- keep-dup 1 -g mm), and peaks overlapping with the blacklisted ones (https://sites.google.com/site/ anshulkundaje/projects/blacklists#TOC-Downloads; https://www.encodeproject.org/annotations/ENCSR636HFF/) were filtered using bedtools-intersect (v2.29.2; -v). The remaining peaks were sorted (bedtools sort) and merged (bedtools merge) into non-overlapping peaks. A consensus peak set was generated by first finding a consensus peakset per condition, and then a global consensus peakset for all conditions using the last ones with bedtools sort and merge. The final counts of aligned reads was done using featureCounts (subread package, v1.6.4).

Only peaks with more than 50 counts in at least one sample were kept. Counts were normalized using the DESeq2 package (v1.28.1). Peaks were annotated using the the ChIPseeker package (v1.24.0). Differentially open peaks were calculated between the controls and mutants (het^+^KO) using the DESeq2 package (v1.28.1). Only peaks that had log2 fold change higher than 1 or lower than −1, and an adjusted p-value lower than 0.1, were considered as significantly differentially open **(Table S1).**

### FAS-regulating transcription factor (TF) and co-transcription factor (co-TF) analysis

FAS-regulating TFs were retrieved from the DoRothEA database (v1.0.1) (21–23) using the expressed genes per cell line, yielding 29 and 30 FAS-regulating TFs in OCI-Ly1 and OCI-Ly8 respectively (**Table S3**), with 21 FAS-regulating TFs expressed in both cell lines. No mutual differentially expressed FAS-regulating TF for OCI-Ly1 and OCI-Ly8 were found.

Only the 21 FAS-regulating TFs which were present in both cell lines were selected for further analysis. Next, a list of all other transcription factors that were expressed in the RNA-seq data was constructed using the DoRothEA database (v1.0.1) (21–23). Only the TFs which were mutually differentially expressed in both OCI-Ly1 and OCI-Ly8 were considered for further analysis, which left 2 down-regulated and 1 up-regulated TFs (**Table S4**). Lastly, the STRING database (v11-0b) (24) was used to construct a network of protein-protein interactions (physical interactions only) between the three mutually differential TFs and the 21 FAS-regulating TFs. Only 8 out of these 21 proteins were predicted to interact with the down-regulated (but not the up-regulated) TFs.

### Statistics

All experiments were done in replicates as indicated. Replicate data is displayed with individual data points, mean and standard deviation (SD). For co-culture experiments, T cells from different donors were used as biological replicates. Data visualization was performed with GraphPad Prism version 6.07 for Windows (GraphPad Software, San Diego, California USA) or in R (v4.2.2) ggplot2 package (v3.4.2). Statistical analysis was performed with R (v4.2.2). The choice of the test, as indicated in the corresponding figure legends, was based on whether the assumptions of the normality of the distributions and homogeneity of the variance between groups were met. These assumptions were tested with the shapiro.test() R function bartlett.test() R function for normally distributed data or flinger.test() R function for non-normally distributed data, respectively. For normally distributed data with homogeneous or non-homogeneous variance, the two-sided t-test (t.test() R function) or the Welch test (t.test() R function with the ’var.equaĺ argument set to FALSE) was used, respectively. For non-normally distributed data with homogeneous variance, the Mann-Whitney U-test (wilcox_test() R function from coin package (v1.4-3)) was used. In case of multiple testing, Bonferroni correction was applied, as indicated in the figure legends. Analysis of RNA and ATAC-sequencing was done with R (v4.2.2) DESeq2 package (v1.38.1) and is described in detail in the respective sections above. For luciferase reporter assay (**Fig 4B**), as well as dose-increasing FASLG-induced apoptosis (**Fig 5B**) testing, linear regression on square root transformed values with lm() R function, with Bonferroni adjustment if needed, was used.

### Cartoon representations

All cartoon representations were created with BioRender.com.

## Acknowledgements

We acknowledge the iFlow Core Facility of the university hospital Munich (INST 409/225-1 FUGG) for assistance with cell sorting. We thank Xavier Pastor Hostench and Dr. Matthias Heinig for the alignment and peak-calling of the ATAC-seq data. We acknowledge Franziska Hackbarth for her laboratory assistance and Miriam Abele for her mass spectrometric support at the BayBioMS. We acknowledge the Core Facility Statistical Consulting of Helmholtz Zentrum Munich, and in particular Dr. Marina Jimenez Munoz, Dr. Gregor Miller, and Prof. Dr. Christian Fuchs, for the consulting on the statistical testing.

## Conflict of Interests statement

M.Sub. receives industry research support from Amgen, BMS/Celgene, Gilead, Janssen, Miltenyi Biotec, Morphosys, Novartis, Roche, Seattle Genetics and Takeda. She serves as a consultant/advisor to AvenCell, CDR-Life, Ichnos Sciences, Incyte Biosciences, Janssen, Miltenyi Biotec, Molecular Partners, Novartis, Pfizer and Takeda and serves on the speakers’ bureau at Amgen, AstraZeneca, BMS/Celgene, Gilead, GSK, Janssen, Novartis, Pfizer, Roche and Takeda. All other authors declare no relevant competing interests.

## Author Contribution Statement

M.A., M.S., M.CT., and O.W. designed the study; M.A. designed and performed experiments, analyzed and interpreted experimental and sequencing data; M.S. analyzed and interpreted sequencing and experimental data; C.D.S. planned, and oversaw co-culture experiments; D.B. designed FASLG-induced apoptosis assay; W.D.K., and J.A.H assisted with interpretation of the data and performed experiments; L.A. assisted with co-culture experiments and interpretation of the data; M.H. performed experiments; V.P. conduced bioinformatics and analysis and interpretation of clinical data; T.F. performed western blot Figure S1; L.W, W.E., S.T., and H.B. performed and oversaw sequencing experiments; M.R. performed multispectral imaging; J.M. performed mass-spectrometry-based proteomic analysis; C.L. oversaw mass-spectrometry-based proteomic analysis; S.B. designed and assisted with CRISPR-Cas9 gene editing procedures; H.L. oversaw CRISPR-Cas9 gene editing procedures; M.S.-S., M.Sub. and M.v.B.-B. helped designing experiments and discussed and interpreted data; M.A., M.S., M.CT., and O.W. wrote the manuscript with input from all authors. O.W. oversaw all experiments and was involved in all aspects of analyzing and interpretation of experimental and clinical data.

## Ethics Statement

All studies herein, including molecular analyses on human material, were covered by IRB approvals (LMU #223-14, LMU #276-14, LMU #056/00, and LMU #539-15 fed).

## Funding Statement

This work was funded by the German Research Foundation (DFG, projects 278529602 - SFB1243 (to O.W. (A11, A12), H.L. (A01) and M.Sub. (A10)), WE 4679/2-1 (to O.W. and M.S.-S.)) and the Wilhelm Sander-Stiftung (2022.093.1 to O.W.). O.W. is supported by the Else Kröner Excellence Fellowship (Else Kröner-Fresenius-Stiftung, 2021_EKES.13) and the Lymphoma Research Foundation (Jaime Peykoff Follicular Lymphoma Initiative). M.Sub. receives funding from the German Research Foundation (DFG, projects 2018.087.1, SFB 338/1 2021 – 452881907 and 451580403). This work was supported by the Impuls-und Vernetzungsfonds of the Helmholtz-Gemeinschaft (grant VH-NG-1219) for M.C.T. and M. S.

## Data Availability Statement

The RNA-sequencing raw data and gene expression matrices have been deposited in the Gene Expression Omnibus (https://www.ncbi.nlm.nih.gov/geo/) under accession number GSE230036. The ATAC-sequencing raw data and peak matrices have been deposited in the Sequence Read Archive (SRA) (https://www.ncbi.nlm.nih.gov/sra) under accession number PRJNA966144. The Proteomic data has been uploaded to ProteomeXchange (https://www.proteomexchange.org) under the identifier PXD041408 and can be accessed on the Panorama web repository server https://panoramaweb.org/2mQCoX.urls. The analysis code is available on Github https://github.com/colomemaria/ARID1A_follicular_lymphoma.

---

**Supplementary Figure 1:**
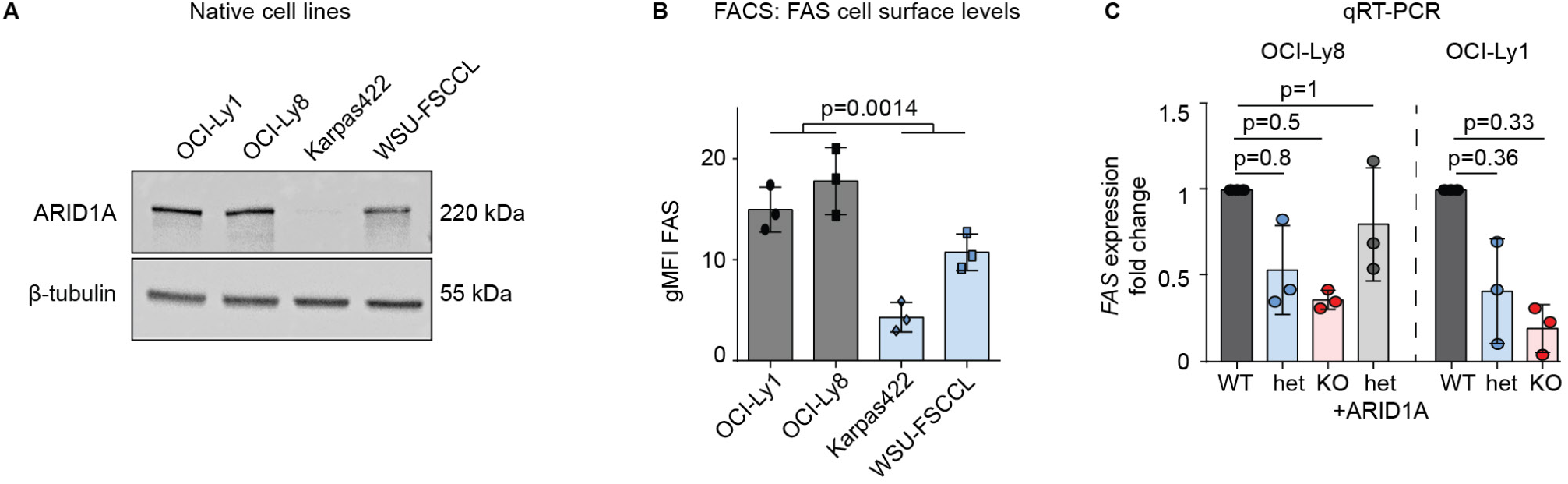
**A** Western blot for ARID1A in *ARID1A*^WT^ OCI-Ly1 and OCI-Ly8 and *ARID1A*^MUT^ Karpas422 and WSU-FSCCL native cells. **B** FAS cell surface expression by FACS on *ARID1A^WT^* (OCI-Ly1 and OCI-Ly8) and *ARID1A^MUT^* (Karpas422 and WSU-FSCCL) native cells. Bar diagram depicting the geometric means of independent replicates (N = 3). *P-value is from two-sided t-test. ARID1A^MUT^ cell lines were tested against ARID1A^WT^ cell lines.* **C** Validation of *FAS* RNA expression by quantitative real-time PCR (TaqMan assay) in OCI-Ly8 and OCI-Ly1 clones (N = 3). *P-values for OCI-Ly1 are from two-sided t-test, for OCI-Ly8 from Mann-Whitney U-test, Bonferroni-adjusted. All groups were tested against WT. Pooled data from biological replicates (N) are represented as mean± SD*.

**Supplementary Figure 2:**
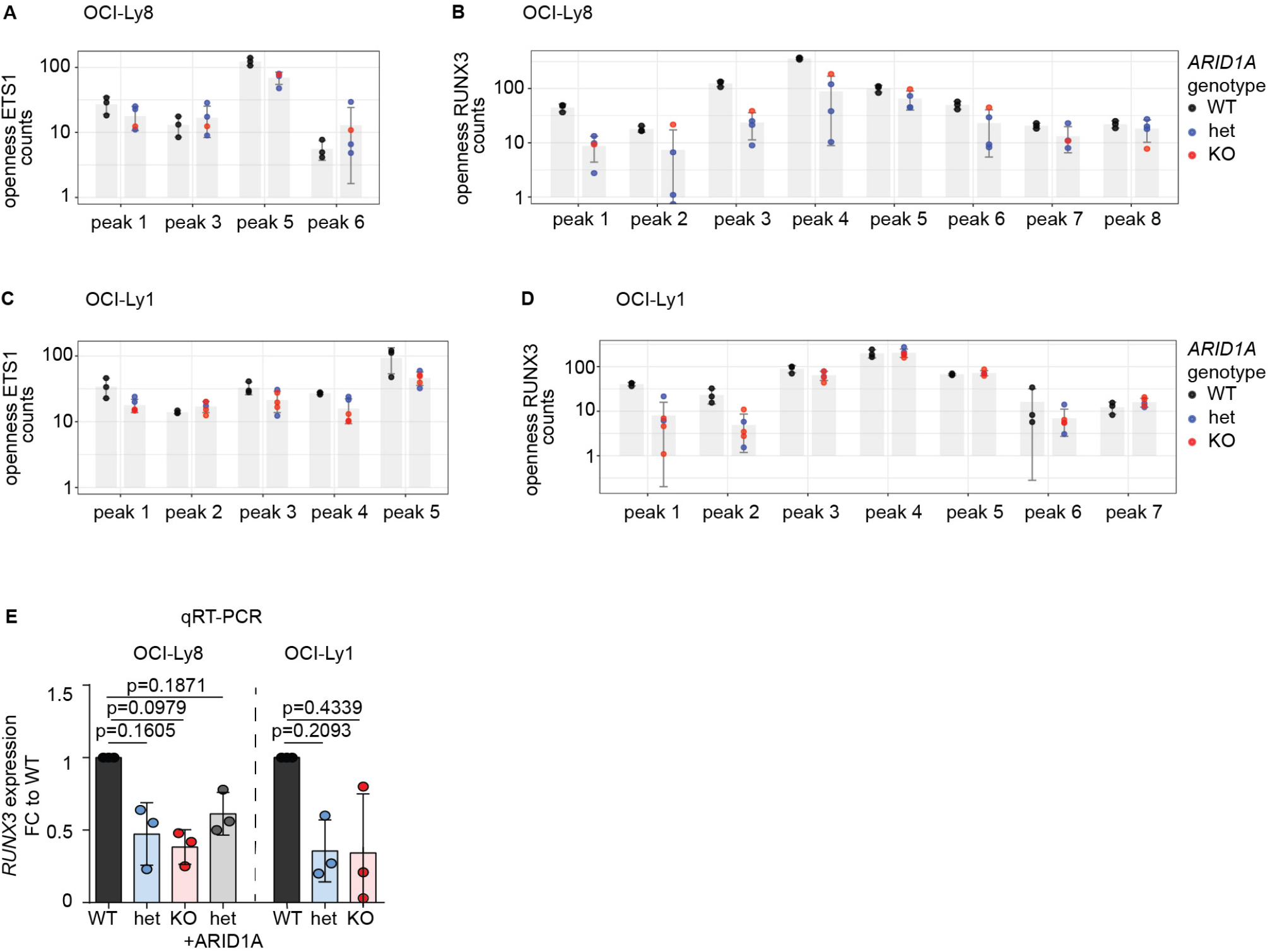
**A** Chromatin openness of ETS1 in OCI-Ly8. **B** Chromatin openness of RUNX3 in OCI-Ly8. **C** Chromatin openness of ETS1 in OCI-Ly1. **D** Chromatin openness of RUNX3 in OCI-Ly1. Black indicates WT, blue indicates *ARID1A*^het^ and red indicates *ARID1A*^hom^. **E** Validation of *FAS* RNA expression by quantitative real-time PCR in OCI-Ly8 and OCI-Ly1 clones (N = 3). *P-values are from the two-sided t-test, Bonferroni-adjusted. All groups were tested against WT. Pooled data from biological replicates (N) are represented as mean± SD*.

**Supplementary Figure 3:**
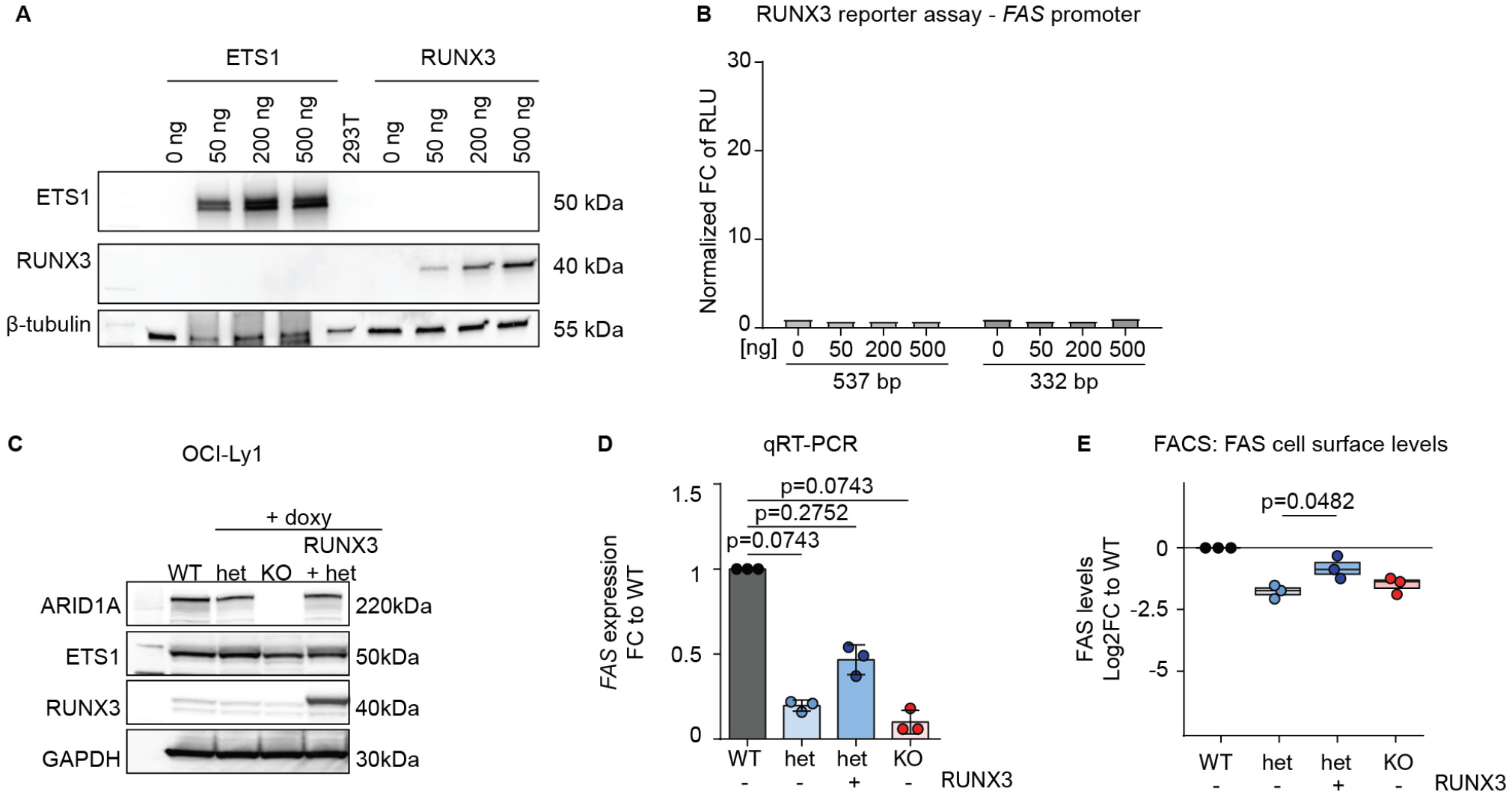
**A** Western blot of ETS1 and RUNX3 in HEK 293T cells after transfection with increasing doses of the respective expression vectors. **B** Luciferase reporter assay, co-transfection of the RUNX3 expression vector and pGL3-FAS constructs (N = 3). **C** Western blot of OCI-Ly1 clones (*ARID1A* WT, het, and KO) with or without stable doxycycline (dox)-induced overexpression of RUNX3. **D** Rescue of *FAS* RNA levels upon RUNX3 overexpression in OCI-Ly1 measured by quantitative real-time PCR (TaqMan assay) (N = 3). *P-values are from Bonferroni-adjusted Mann-Whitney U-test. All groups were tested against WT.* **E** Rescue of FAS cell-surface protein levels upon RUNX3 overexpression in OCI-Ly1 measured by FACS (N = 3). *P-value is from two-sided t-test. Het+RUNX3 was tested against het. Pooled data from biological replicates (N) are represented as mean*± *SD*.

## Notes

https://github.com/colomemaria/ARID1A_follicular_lymphoma

